# Concordance of claudin-18.2 expression in biopsy, resection, and recurrent specimens: implications for zolbetuximab therapy in pancreatic ductal adenocarcinoma

**DOI:** 10.1101/2025.04.11.648482

**Authors:** Daisuke Kyuno, Kazuhiko Yanazume, Akira Saito, Yusuke Ono, Tatsuya Ito, Masafumi Imamura, Makoto Osanai

## Abstract

Claudin-18.2 is a promising therapeutic target for gastrointestinal cancer. However, its expression pattern in pancreatic ductal adenocarcinoma, especially the concordance between biopsy and resection specimens, is unknown. This study aimed to evaluate the consistency of claudin-18.2 positivity across different specimen types using the clinically validated antibody clone 43-14A employed in ongoing zolbetuximab trials. Immunohistochemical analysis for claudin-18 was conducted on 211 resected pancreatic cancer tissues, 133 matched preoperative biopsy samples, and 60 samples from recurrent lesions. Concordance rates were calculated based on a clinically relevant cutoff (≥75% of tumor cells with moderate-to-strong membranous staining). Receiver operating characteristic analysis was used to optimize the biopsy thresholds. Claudin-18.2 positivity was observed in 9.5% of the resection specimens. The concordance rates were 92.5% between biopsy and resection specimens and 83.3% between primary and recurrent lesions. Receiver operating characteristic analysis suggested that a lower cutoff (20%) in biopsy samples achieved 100% sensitivity for detecting positive cases. While overall claudin-18 expression levels were maintained in most recurrent lesions, decreased expression was frequently observed in cases of local recurrence and liver metastasis. Despite intrinsic heterogeneity and limited biopsy yield in pancreatic cancer, claudin-18 expression in small biopsy samples, assessed using the clinical trial-validated clone 43-14A, strongly correlated with that in resection specimens. These results suggest that claudin-18.2 is a clinically stable biomarker for zolbetuximab therapy in patients with pancreatic ductal adenocarcinoma.

**Graphical abstract:** 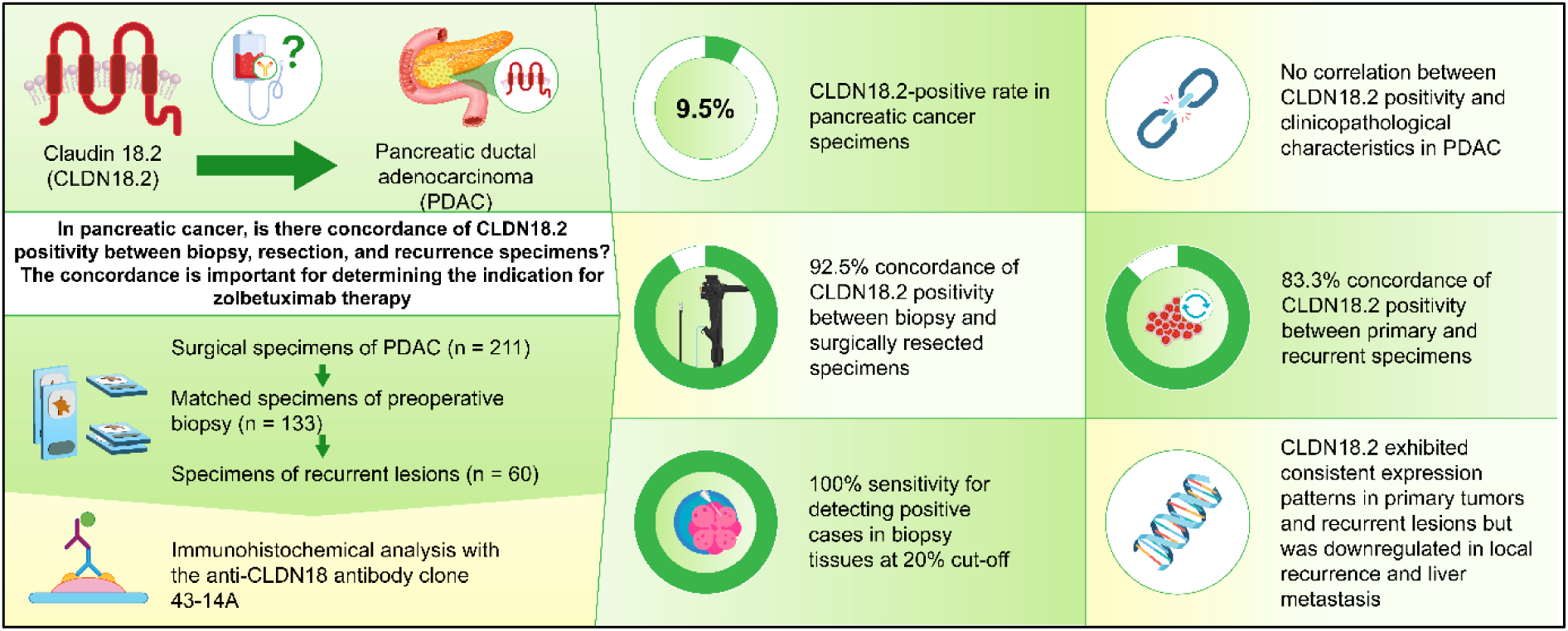

**Core Tips:**

1. This study demonstrated a high concordance (92.5%) of CLDN18.2 positivity between preoperative biopsy and resected pancreatic ductal adenocarcinoma specimens, using the clinical trial-validated antibody clone 43-14A.
2. CLDN18.2 expression in recurrent lesions generally remained consistent with that in the corresponding primary tumor, although a reduction in expression levels was observed in local recurrence and liver metastases.
3. These findings support the clinical utility of biopsy-based CLDN18.2 evaluation for selecting patients with pancreatic ductal adenocarcinoma who may benefit from zolbetuximab.

## Introduction

Claudin-18.2 (CLDN18.2), a tight junction protein predominantly expressed in gastric tissues, has been identified as a promising therapeutic target for various gastrointestinal malignancies, including gastric and pancreatic cancers. The monoclonal antibody zolbetuximab, which specifically targets CLDN18.2, has demonstrated efficacy in improving the survival of patients with unresectable advanced gastric and gastroesophageal junction adenocarcinomas in the phase 3 clinical trials GLOW and SPOTLIGHT.^1,2^ These trials showed significantly prolonged median progression-free survival and overall survival in patients with CLDN18.2-positive cancer. Given these positive outcomes, zolbetuximab is anticipated to be a key component of future treatment protocols for pancreatic ductal adenocarcinoma (PDAC).

In an ongoing phase 2 clinical trial for metastatic pancreatic cancer (ClinicalTrials.gov ID: NCT03816163), patients treated with zolbetuximab are selected based on the positive expression of CLDN18.2, as determined by immunohistochemical staining. The criteria for CLDN18.2 positivity, consistent with those of the GLOW and SPOTLIGHT trials, require moderate-to-strong membrane staining of CLDN18.2 in at least 75% of the cancer cells.^1–3^ However, previous immunohistochemical analyses of resected gastric and pancreatic cancer specimens revealed heterogeneous CLDN18 staining intensity across tissue regions within the same samples.^4–7^ This variability raises concerns regarding the accuracy of CLDN18.2 expression assessments based on biopsy specimens, which may not accurately represent the staining pattern observed in the entire resected tumor.

Recent studies on gastric cancer tissues reported substantial heterogeneity in CLDN18.2 expression across tumor regions, indicating that the choice of biopsy site influences the assessment of CLDN18.2 expression.^7,8^ This finding suggests the need for similar studies on PDAC to ensure the reliability of biopsy-based evaluations, as discrepancies between biopsy and resection specimens may affect the accuracy of patient selection for zolbetuximab therapy. Furthermore, for patients with unresectable PDAC, which account for more than 75% of cases,^9,10^ therapeutic regimen will depend on the CLDN18.2 positivity in limited biopsy tissue, understanding the relationship between CLDN18.2 expression in biopsy and resection specimens is critical. The accurate assessment of biopsy specimens is essential to determine the eligibility for zolbetuximab treatment, making this area of research highly relevant for optimizing therapeutic outcomes.

Additionally, examining the concordance of CLDN18.2 expression between primary and recurrent PDAC tissues is crucial for determining the applicability of zolbetuximab in recurrent cases. Biopsies from metastatic PDAC lesions, typically identified using imaging modalities,^11^ are not always available. Thus, understanding the relationship between CLDN18.2 expression in primary and recurrent sites may provide valuable insights for clinical decision-making, making the findings of the present study particularly interesting for the management of patients with recurrent disease.

To date, no study has systematically compared the concordance of CLDN18.2 positivity between biopsy and resected PDAC specimens or between primary and recurrent PDAC specimens using the anti-CLDN18 antibody clone 43-14A, the same clone used in clinical trials.^1,2^ Assessing the concordance rates may optimize patient selection for zolbetuximab therapy, thus providing crucial data for improving cancer treatment strategies.

## Materials and methods

### Sample collection

Specimens from 222 consecutive patients with pancreatic cancer who underwent pancreatomy between January 2011 and December 2020 were retrieved from the pathology files of Sapporo Medical University Hospital. Follow-up data were obtained from hospital records. The inclusion criteria for resection specimens were that pancreatic cancer was pathologically confirmed in these specimens. The pancreatic cancers included PDAC, acinar cell carcinoma, carcinoma derived from intraductal papillary mucinous neoplasia, adenosquamous carcinoma, and anaplastic carcinoma.

Patients with pathological complete response after neoadjuvant therapy (n = 7), those diagnosed with pancreatic intraepithelial neoplasia (PanIN) only (n = 3), and those with specimens containing only a small number of cancer cells (n = 1) were excluded. Thus, specimens from 211 patients were analyzed.

Preoperative biopsy tissues were obtained using endoscopic ultrasound-guided fine-needle aspiration (EUS-FNA) in all cases. The inclusion criteria for preoperative biopsy specimens were as follows: (1) pancreatic cancer was pathologically confirmed in these specimens and (2) the specimens contained a sufficient number of cancer cells for quantitative evaluation of immunostaining. Specimens diagnosed with suspected pancreatic cancer were excluded from the study. Biopsy specimens that met the inclusion criteria were obtained from 133 patients.

Specimens from recurrent lesions were obtained via resection or needle biopsy. The inclusion criteria for such samples were that tissue from recurrent lesions contained (1) pathologically confirmed metastatic carcinoma cells and (2) a sufficient number of cancer cells for quantitative evaluation of immunostaining. Thus, samples from 60 recurrent lesions in 53 patients were collected, as seven patients had recurrent lesions in two different organs.

The 211 cases were staged according to the UICC staging system (8th edition).^12^ Tumor resectability status was classified according to the NCCN and Japanese criteria.^13,14^ Preoperative CEA and CA19–9 levels were measured within four weeks prior to surgery. Postoperative surveillance was performed monthly during the first postoperative year and every one to three months thereafter. Recurrence was diagnosed based on radiological findings and confirmed through biopsy if possible.

### Immunohistochemical analysis of specimens

Immunohistochemistry was performed using the anti-CLDN18 antibody clone 43-14A (1:5000; Abcam, ab314690, Cambridge, UK), which was previously used to determine CLDN18.2 positivity in gastric cancer^3^. In the present study, tissue sections were deparaffinized in xylene, rehydrated using a graded series of ethanol and phosphate-buffered saline. After antigen retrieval by microwave heating (95°C for 30 min) in 10 mmol/L Tris/1 mmol/L EDTA buffer, the sections were incubated with 3% H_2_O_2_ for 10 min to block endogenous peroxidase activity, followed by 4% skim milk as a blocking buffer for 30 min, and then with antibodies diluted in Dako REAL™ Antibody Diluent (Agilent, Santa Clara, CA, USA) overnight at 4°C. The sections were then incubated with the Dako REAL™ EnVision™ HRP rabbit/mouse for 30 min at room temperature, and color was developed using the Dako REAL™ EnVision™ DAB plus CHROMOGEN and substrate buffer for 10 min, according to the manufacturer’s instructions. The slides were then counterstained with hematoxylin.

As negative controls, adjacent non-neoplastic regions were examined as normal tissues. As a positive control, normal gastric mucosal tissue was stained simultaneously with PDAC tissue (Supplemental Figure 1A). The staining intensity of the surgical specimens was scored as follows: 0, no reactivity at the cellular membrane; 1+, weak reactivity; 2+, moderate reactivity; and 3+, strong reactivity, as in gastric cancer (Supplemental Figure 1B).^8,15^ The proportion of the stained area was scored between 0 and 100 in increments of 5. The histological scores (H-scores) of PDAC, low-grade PanIN, and normal ducts were obtained by multiplying the proportion and intensity scores, therefore ranging between 0 and 300. Because all high-grade PanINs were difficult to differentiate from the intraductal spread of cancer, the H-score of high-grade PanIN alone was not analyzed. CLDN18 staining intensity did not correlate with the year of the surgical tissue collection (Supplemental Figure 1C). CLDN18.2 positivity was defined as ≥75% of areas with moderate-to-strong staining intensity; the same criterion was employed in previous clinical trials.^1–3^ The stained areas were evaluated as the percentage of the number of stained cancerous glandular ducts, even partially, divided by the total number of cancerous glandular ducts.

Two pathologists (DK and MO) were blinded to the clinical data when evaluating the slides. Discordant cases were discussed, and a consensus on the staining intensity and area was reached.

### Statistical analysis

The measured values are presented as mean ± standard deviation. Data were analyzed and compared using Fisher’s exact test and the Mann–Whitney U test. The correlation between two variables was tested using Spearman’s correlation coefficient. The difference between two paired data was tested using the Wilcoxon signed-rank sum test. Survival rates were calculated using the Kaplan–Meier method and compared using the log-rank test and a Cox proportional hazard model to estimate hazard ratios (HRs) and 95% confidence intervals (CIs). To detect CLDN18.2 true-positive cases using biopsy specimens, receiver operating characteristic (ROC) analysis was conducted to establish a suitable cutoff proportion for the moderate-to-strong staining area of CLDN18 in biopsy samples.

Youden Index is used to identify the point on an ROC curve where both sensitivity and specificity reach their peak values. This point represents the highest vertical distance from the diagonal line, indicating no discrimination on the ROC curve.^16^ Statistical significance was set at P < 0.05. All statistical analyses were performed using BellCurve for Excel version 4.08 and EZR software version 1.68.^17^

## Results

### Expression profiles of claudin-18 in surgical specimens of PDAC, PanIN, and normal ducts

We examined the expression and distribution of CLDN18 in PDAC, PanIN, and normal pancreatic ducts of resected PDAC specimens using immunohistochemical staining with the anti-CLDN18 antibody 43-14A. Strong CLDN18 expression was detected in PanIN, moderate expression in PDAC, and almost no expression in normal ducts (Figure 1A–C). CLDN18 was located on the plasma membrane of all tissue types. In the continuous area between low- and high-grade PanIN, CLDN18 staining delignated the low- and high-grade areas with a clear boundary (Figure 1D). In addition, moderate-to-strong levels of CLDN18 staining were observed in acinar-to-ductal metaplasia (Figure 1E). This metaplasia was excluded from the evaluation of the staining intensity and area. Even within the same tissue, the CLDN18 staining properties of cancer cells were heterogeneous (Figure 1F).

**Figure 1.**
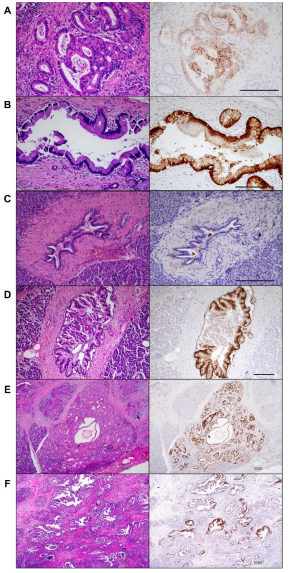
Representative images of CLDN18 immunohistochemical stainings. The left panels show images of hematoxylin and eosin stainings, and the right panels show CLDN18 immunohistochemistry images of PDAC (A), low-grade PanIN (B), normal pancreatic duct (C), low- and high-grade PanIN (D), acinar-to-ductal metaplasia in the pancreas (E), and low-power fields of PDAC (F). Scale bar: 200 µm. CLDN18, claudin-18; PDAC, pancreatic ductal adenocarcinoma; PanIN, pancreatic intraepithelial neoplasia.

The intensity of CLDN18 expression was analyzed using immunohistochemistry in 211 resected pancreatic cancer tissues. The H-score of CLDN18 was significantly higher in PanIN than in PDAC, and CLDN18 was not expressed in almost all normal pancreatic ducts (P < 0.001 between tissue types; Figure 2A). CLDN18 expression was not detected in adenosquamous (n = 3), anaplastic (n = 1), or acinar cell carcinomas (n = 2). One case of invasive intraductal papillary carcinoma showed 20% of moderate-to-strong CLDN18 staining, and three cases did not show any moderate-to-strong CLDN18 staining. In total, 172 cases (72.0%) were positive for any CLDN18 staining, and the intensity of CLDN18 staining was heterogeneous within the same pancreatic cancer specimen (Figure 2B). CLDN18 was not expressed in 59 cases (28.0%), and 56 cases (26.5%) showed such heterogeneity that both no-to-weak and moderate-to-strong expression regions represented more than 25% in the same specimen. In the SPOTLIGHT and GLOW studies, CLDN18.2 positivity was defined as ≥75% of tumor cells demonstrating moderate-to-strong membranous CLDN18 staining ^1–3^. In the current study, the prevalence of CLDN18.2 positivity was 9.5% (20/211) among all screened patients (Figure 2C).

**Figure 2.**
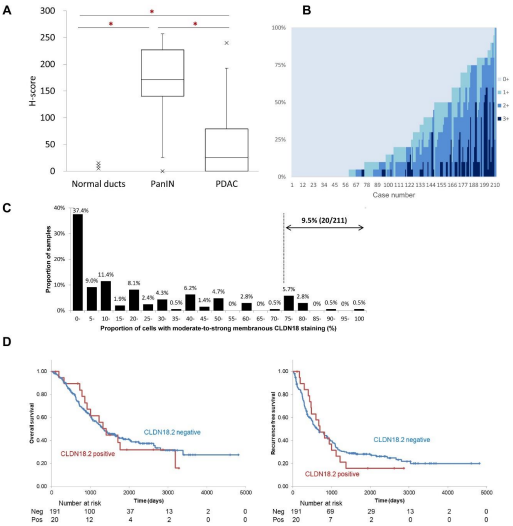
CLDN18 expression in PDAC. (A) H-score of CLDN18 in normal ducts, PanIN, and PDAC. Statistical significance was determined using the Mann–Whitney U test. *P < 0.001. (B) Overall expression intensity of CLDN18 in PDAC tissues. (C) Distribution of CLDN18 staining in all screened patients. The numbers above each bar indicate the proportion of samples within each bin.

Using information from our hospital patient database, we performed a survival analysis of patients with PDAC regarding CLDN18.2 positivity. No significant correlation was observed between CLDN18.2 positivity and any clinicopathological feature (Supplemental Table 1). CLDN18.2 positivity was not associated with recurrence-free survival (HR: 1.05, 95% CI: 0.63–1.77) and overall survival (HR: 1.04, 95% CI: 0.59–1.86; Figure 2D).

The dotted line indicates the cutoff for CLDN18.2 positivity (≥75% of tumor cells with moderate-to-strong membranous CLDN18 staining). (D) Kaplan–Meier curves for overall survival (left) and recurrence-free survival (right) of patients with PDAC grouped by CLDN18.2 positivity. CLDN18, claudin-18; H-score, histological score; PDAC, pancreatic ductal adenocarcinoma.

### Comparison between preoperative biopsied and resection specimens

CLDN18 expression was compared in 133 cases in which both preoperative biopsy and resection specimens of the primary tumor were available for analysis. The proportions of samples with moderate-to-strong membrane staining were correlated between biopsy and resection specimens (Spearman’s rank correlation coefficient: 0.500, P < 0.001; Figure 3A). The concordance rate of CLDN18.2 positivity between these specimens was 92.5% (123/133; P < 0.001, Fisher’s test) at a 75% cutoff value (Table 1). The concordance of CLDN18.2 positivity did not correlate with any clinicopathological features (Supplemental Table 2), and the presence or absence of neoadjuvant treatment did not correlate with the difference in the proportion of moderate-to-strong CLDN18 staining areas between biopsy and resection specimens (P = 0.760, Mann–Whitney U test).

**Table 1.** Comparison of CLDN18.2 positivity between matched biopsy and resection specimens.

|  | Characteristic | Resection |  | Total |
| --- | --- | --- | --- | --- |
| | | $\geq 75\%$ | $< 75\%$ | |
| Biopsy | $\geq 75\%$ | 6 | 5 | 11 |
| | $< 75\%$ | 5 | 117 | 122 |
| Total |  | 11 | 122 | 133 |

**Figure 3.**
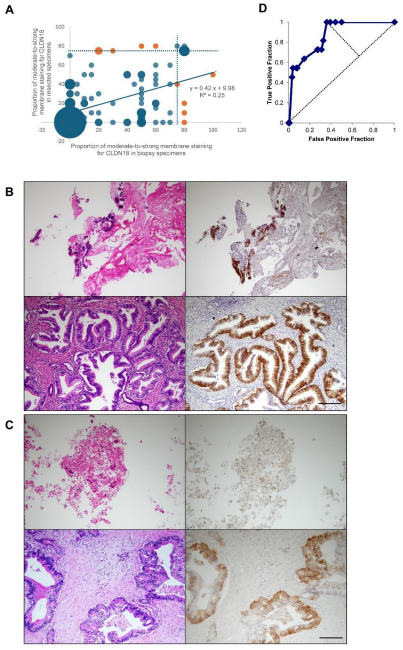
Correlation of CLDN18.2 positivity between biopsy and resection specimens. (A) Bubble chart showing the proportions of moderate-to-strong membranous staining for CLDN18 in biopsy and resection specimens. The bubble size indicates the number of cases. Dotted lines indicate 75%. Orange bubbles represent cases in which the CLDN18.2 positivity of a biopsy specimen did not match that of the corresponding resection specimen. (B and C) Representative images of cases in which the positivity of biopsy and resection specimens matched (B) and did not match (C). The left panels show images of hematoxylin and eosin staining, and the right panels show CLDN18 immunohistochemistry images of PDAC. (D) ROC curve of the proportion of moderate-to-strong CLDN18 staining of biopsy specimens that is indicative for the CLDN18.2 positivity of resection specimens. CLDN18.2, claudin-18.2; PDAC, pancreatic ductal adenocarcinoma; ROC, receiver operating characteristic.

The sensitivity of biopsied specimens was not high at 54.6% (6/11); five unmatched cases showed negative results in the biopsied specimens but CLDN18.2 positivity in the resection specimens (Figure 3B and C). To minimize false negatives, ROC analysis was performed to determine the percentage of moderate-to-strong membrane staining of biopsy tissues that would indicate CLDN18.2 positivity in resection specimens (Figure 3D). The area under the curve was 0.866 (95% CI: 0.774– 0.958, P < 0.001), and 20% of moderate-to-strong CLDN18 staining showed the highest Youden index (0.639). At a 20% cutoff in biopsy tissues, the sensitivity was 100% (11/11), and the concordance rate was 66.9% (89/133; P < 0.001, Fisher’s test; Table 2).

**Table 2.**
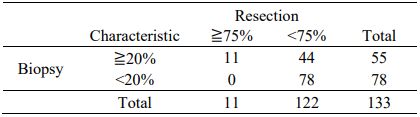
Comparison of CLDN18.2 positivity between matched biopsy at a 20% cutoff and resection specimens at a 75% cutoff.

### Comparison between primary and recurrent specimens

CLDN18 expression was compared in 53 patients and 60 samples in which both primary tumors and recurrent tissues were available for analysis. The recurrent sites of the tissues used in this analysis were local sites (remnant pancreas or tumor beds; n = 15), the liver (n = 14), lungs (n = 18), peritoneum (n = 6), and others (para-aortic lymph nodes and tumor cells in pleural effusion). The proportions of moderate-to-strong membrane stainings were correlated between primary and recurrent specimens (Spearman’s rank correlation coefficient: 0.396, P = 0.002; Figure 4A). The concordance rate of CLDN18.2 positivity between these specimens was 83.3% (50/60; P = 0.062, Fisher’s test) at a 75% cutoff (Table 3). The concordance of CLDN18.2 positivity did not correlate with any clinicopathological features, including the presence or absence of adjuvant treatment (Supplemental Table 3). In many cases of local recurrence and liver metastasis, the proportions of moderate-to-strong CLDN18 stainings were reduced in recurrent lesions compared to those in the corresponding primary lesions (P = 0.029 for local recurrence, P = 0.175 for liver metastasis, Wilcoxon signed-rank sum test; Figure 4B–D). The overall survival rate was not correlated with CLDN18.2 positivity of recurrent specimens (HR: 1.144, 95% CI: 0.439–2.984) and the concordance between primary and recurrent specimens (HR: 1.358, 95% CI: 0.477–3.870).

**Figure 4.**
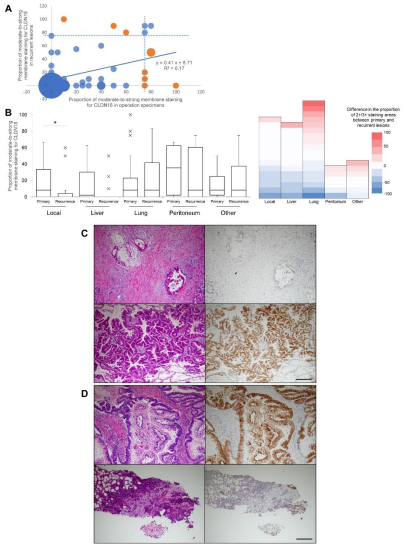
Correlation of CLDN18.2 positivity between primary and recurrent specimens. (A) Bubble chart showing the proportions of moderate-to-strong membranous staining for CLDN18 in primary and recurrent specimens. The bubble size indicates the number of cases. Dotted lines indicate 75%. Orange bubbles represent cases in which the CLDN18.2 positivity of a primary specimen did not match that of the corresponding recurrent specimen. (B) Proportion of moderate-to-strong membrane staining for CLDN18 in primary and recurrent specimens. Statistical significance was determined using the Wilcoxon signed-rank sum test. *P = 0.029. The heatmap shows the difference in the proportion of moderate-to-strong staining areas between primary and recurrent lesions (proportion of recurrent lesions minus the proportion of primary lesions). (C and D) Representative images of cases in which the positivity of the primary and recurrent specimens did not match. The left panels show images of hematoxylin and eosin stainings, and the right panels show CLDN18 immunohistochemistry images. (C) CLDN18-negative staining in primary PDAC (upper panels) and positive staining in lung metastasis (lower panels). (D) CLDN18-positive staining in primary PDAC (upper panels) and negative staining in liver metastasis (lower panels). CLDN18.2, claudin-18.2; PDAC, pancreatic ductal adenocarcinoma.

**Table 3.** Comparison of CLDN18.2 positivity between matched primary and recurrent specimens.

|  |  | Recurrence |  | Total |
| --- | --- | --- | --- | --- |
| Characteristic | | $\geq 75\%$ | $< 75\%$ | |
| Primary | $\geq 75\%$ | 3 | 6 | 9 |
| | $< 75\%$ | 4 | 47 | 51 |
| Total |  | 7 | 53 | 60 |

## Discussion

This study demonstrated a significant concordance in CLDN18.2 positivity between preoperative biopsy and resected PDAC specimens, as well as between primary and recurrent PDAC lesions. Specifically, we found a high concordance rate of 92.5% at the clinically relevant 75% cutoff in biopsy–resection comparisons and 83.3% concordance in primary–recurrence comparisons. These findings suggest that the immunohistochemical evaluation of biopsy samples can reliably reflect CLDN18.2 expression in the entire tumor and can therefore serve as a robust basis for patient selection for zolbetuximab therapy. This study assessed the consistency of CLDN18.2 expression across both biopsy and resection samples and across primary and recurrent PDAC, providing novel insights into the clinical applicability of CLDN18.2-targeted treatment strategies.

The high concordance rate (92.5%) of CLDN18.2 positivity between biopsy and resected PDAC specimens highlights the potential reliability of preoperative biopsy to facilitate patient stratification for CLDN18.2-targeted therapies. This finding is clinically significant because treatment decisions for unresectable PDAC are often based solely on biopsy samples. In gastric cancer, the concordance rate of CLDN18.2 positivity in biopsy and resection specimens was 81.3% at the 75% cutoff, and the superficial area of the tumor tended to have a higher CLDN18-positive rate than the invasive front.^7^ Previous studies on gastric cancer have reported substantial intratumor heterogeneity in CLDN18.2 expression, raising concerns about false-negative results due to sampling bias.^4,5^ Likewise, pancreatic cancer exhibits considerable intratumoral heterogeneity, extensive stromal fibrosis, and microenvironmental complexity, potentially exceeding that of gastric cancer in terms of sampling variability.^18^ In light of these characteristics, the observed high concordance rate is particularly noteworthy.

One possible explanation for the high concordance is the relatively low overall prevalence of CLDN18.2 positivity in PDAC, as shown in this study (9.5% positive). The CLDN18.2 positivity rate for pancreatic cancer observed in this study is lower than the rate of 38.4% for gastric cancer^3^ but similar to the rate of 13.1% for cholangiocarcinoma.^19^ When most cases are negative in both biopsy and resection specimens, the apparent concordance rate may be inflated. Nevertheless, ROC analysis demonstrated that using a lower cutoff value (20%) in biopsies achieved 100% sensitivity in identifying patients who would be CLDN18.2-positive in resection specimens. Although caution should be exercised in increasing the number of false positives owing to changes in cutoff values, this finding suggests that threshold adjustment in biopsy evaluation may help mitigate the false-negative risk while maintaining adequate concordance for clinical decision-making.

The concordance of CLDN18.2 positivity between primary and recurrent PDAC lesions was slightly lower (83.3%) than that observed between biopsy and resection specimens. The concordance rates of CLDN18.2 positivity between primary and recurrent lesions differ depending on the tissue type and primary site of the cancer. For gastric cancer, the concordance rate between the primary tumor and its metastases ranged from 74.8% to 81.5%,^5,7,8^ whereas for ovarian cancer, it was 58% in mucinous tubo-ovarian carcinoma depending on the histological subtype.^20^ Although this concordance is relatively high, it should be interpreted with caution. In some cases, particularly those with local recurrence or liver metastasis, the proportion of CLDN18.2-positive cells was reduced in recurrent lesions compared to that in primary tumors. Similarly, decreased concordance rates in liver metastases have been observed in gastric cancer.^5^ This observation may reflect clonal selection or phenotypic shifts associated with tumor progression, therapeutic interventions, or changes in the tumor microenvironment during metastasis. Despite these changes, our findings suggest that CLDN18.2 expression in primary tumors may, in most cases, serve as a surrogate for its expression in recurrent sites, which is clinically valuable given the limited feasibility of re-biopsy in recurrent PDAC. These results support the potential utility of initial CLDN18.2 assessment in guiding treatment decisions for both primary and recurrent diseases, although confirmatory studies with larger cohorts and more metastatic samples are warranted.

The clinical implications of this study extend beyond patient selection for zolbetuximab therapy. The reliable detection of CLDN18.2 expression in preoperative biopsy specimens and moderate concordance with recurrent lesions suggest that CLDN18.2 may serve as a stable biomarker throughout disease progression in a subset of PDAC cases. As molecular targeted therapy becomes increasingly integrated into the management of pancreatic cancer,^21,22^ the accurate and minimally invasive evaluation of biomarkers will be essential for identifying candidates for targeted therapies. Importantly, biopsy sampling for pancreatic cancer is largely limited to EUS-FNA. In gastric cancer, biopsy by forceps under direct endoscopic view is possible, and multiple biopsies increase the sensitivity of CLDN18.2 positivity.^8^ However, in pancreatic cancer, EUS-FNA can only be performed a limited number of times because of its invasive nature and often yields small samples with few viable tumor cells because a large fraction of the tissue was composed of blood cells and fibrotic tissue around the cancer cells. In this context, the high concordance between CLDN18.2 positivity in biopsy and resection specimens, as observed in the present study, is particularly valuable, as it demonstrates that even limited biopsy samples can provide reliable information for clinical decision-making in PDAC.

Although several studies have investigated CLDN18.2 expression in pancreatic cancer, only three— including the present study—have employed the clone 43-14A antibody, which is the same clone used in ongoing clinical trials.^23,24^ Among these, our study included the largest number of PDAC specimens to date. Notably, this is the only study to systematically and quantitatively compare CLDN18.2 expression across biopsy, resection, and recurrent specimens. These methodological strengths enhance the clinical relevance and generalizability of our findings to the general population.

This study has several limitations. First, the retrospective nature and single-center design may limit the generalizability of our findings. Second, the methods for CLDN18 immunostaining used in this study were different from those in gastric cancer clinical trials. In the trials, CLDN18 was stained with the VENTANA CLDN18 (43-14A) RxDx Assay (VMSI/Roche) on a BenchMark ULTRA instrument (VMSI/Roche). Consequently, we assessed the staining intensity in both positive and negative controls each time immunostaining was performed. Third, the relatively low proportion of CLDN18.2-positive cases may have contributed to an overestimation of concordance, owing to the high number of true-negative pairs. Forth, the number of recurrent specimens, particularly from metastatic sites, was limited. Finally, the influence of treatment history on CLDN18.2 expression could not be fully evaluated. These limitations might be related to patient cohort bias, including race and treatment modality, which may change the percentage of CLDN18.2-positive patients. In this case, the concordance rate of CLDN18.2 positivity among biopsy, resection, and recurrent specimens and the cutoff values in biopsied tissues would also change. Future multicenter studies involving larger cohorts, especially with more CLDN18.2-positive cases and diverse metastatic sites, are warranted to validate and refine our findings. In addition, longitudinal monitoring of CLDN18.2 expression before and after treatment may help to clarify its temporal stability and predictive value. Despite these limitations, the present study provides a foundation for incorporating CLDN18.2 testing into the diagnostic and therapeutic workflows for pancreatic cancer and highlights its potential role as a reliable biomarker in both localized and recurrent disease contexts.

## Conclusion

This study demonstrated that CLDN18.2 expression in biopsy specimens of patients with PDAC correlated strongly with that in resected tissues and, to a lesser extent, with that in recurrent lesions. Despite the intrinsic heterogeneity and limited tissue availability in PDAC, the immunohistochemical evaluation of CLDN18 in preoperative biopsy samples appears to provide reliable clinical information. These findings support the feasibility of using biopsy-based CLDN18.2 assessment to identify candidate patients for zolbetuximab therapy, even in the context of recurrent disease, where repeat biopsies are often impractical. While further validation in larger prospective studies is warranted, our results provide valuable evidence supporting the integration of CLDN18.2 testing into routine diagnostic and therapeutic decision-making workflows for PDAC.

Given the dismal prognosis of pancreatic cancer and the emerging potential of zolbetuximab to improve patient survival, we hope that the findings of this study will help facilitate appropriate patient selection and ultimately contribute to better clinical outcomes for individuals affected by this devastating disease.

## Acknowledgments

We would like to thank Yui Kawami and Fuminori Daimon for technical assistance with the experiments, Reona Okumura and Hinae Asano MD-PhD students at Sapporo Medical University for sampling and statistical analysis, and Editage (http://www.editage.com) for editing and reviewing this manuscript for English language.

## Data availability statement

The other datasets generated during and/or analyzed during the current study are available from the corresponding author upon reasonable request.

## Ethics approval

This study was approved by the Institutional Review Board of Sapporo Medical University (IRB study number 292-68). The study was approved on August 25, 2017. The need for informed consent was waived by the Institutional Review Board of Sapporo Medical University because of the retrospective nature of the study.

## Author Contributions

Conceptualization, D.K.; methodology, D.K; validation, D.K., K.Y., M.O.; formal analysis, D.K., K.Y.; investigation, D.K., K.Y., A.S., Y.O., M.O.; resources, D.K., T.I., M.I.; data curation, D.K., K.Y.; writing—original draft preparation, D.K.; writing—review and editing, D.K.; visualization, D.K., K.Y.; supervision, D.K.; project administration, D.K.; funding acquisition, D.K., M.O. All authors have read and agreed to the published version of the manuscript.

