## Supplemental Tables and Figure for "Concordance of claudin-18.2 expression in biopsy, resection, and recurrent specimens: implications for zolbetuximab therapy in pancreatic ductal adenocarcinoma"

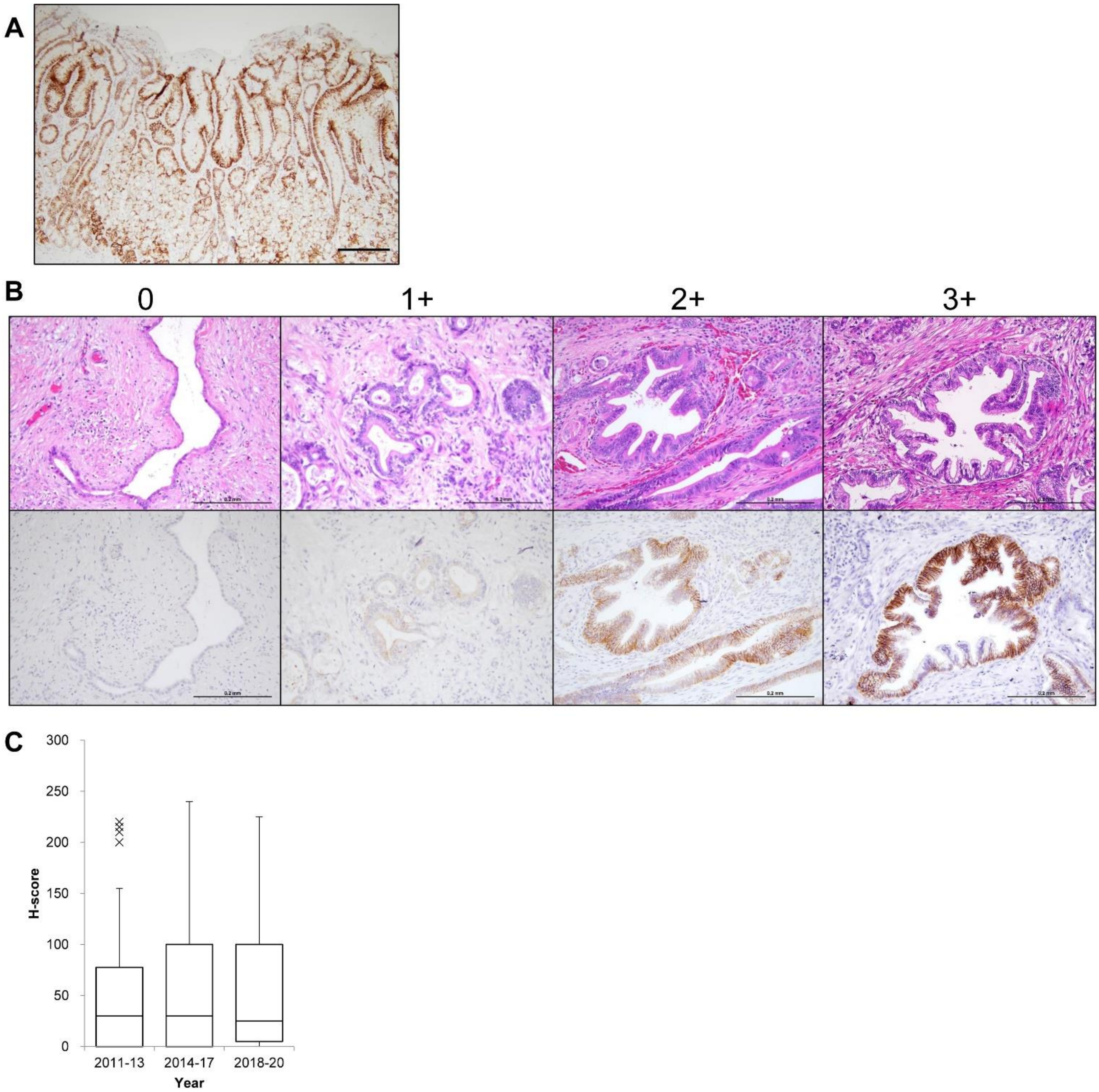


**Supplemental Figure 1**. Representative images of CLDN18 immunohistochemistry. (A) Immunohistochemical staining for CLDN18 in healthy gastric mucosa. (B) Staining intensity of CLDN18 immunohistochemistry. 0, no reactivity in the cellular membrane; 1+, weak reactivity; 2+, moderate reactivity; and 3+, strong reactivity, as in gastric cancer. (C) H-score of CLDN18 according to the year of surgical tissue collection. CLDN18, claudin-18; H-score, histological score.

| Variables | Cldn18 negative | Cldn18 positive | p-value |
|---|---|---|---|
| Overall | 191 (90.5) | 20 (9.5) | |
| Age | | | 0.819 |
| ≤70 | 101 (47.9) | 10 (4.7) | |
| >70 | 90 (42.7) | 10 (4.7) | |
| Sex | | | 0.640 |
| Female | 91 (43.1) | 11 (5.2) | |
| Male | 100 (47.4) | 9 (4.3) | |
| ASA-PS classification system | | | 0.440 |
| 0-2 | 155 (73.5) | 15 (7.1) | |
| 3 | 19 (9.0) | 3 (1.4) | |
| BMI | | | 0.765 |
| ≤25 | 139 (65.9) | 14 (6.6) | |
| >25 | 35 (16.6) | 4 (1.9) | |
| Tumor location | | | 0.083 |
| Head | 133 (63.0) | 10 (4.7) | |
| Body/Tail | 58 (27.5) | 10 (4.7) | |
| Resectability | | | 0.811 |
| R | 133 (63.0) | 63 (29.9) | |
| BR/UR | 14 (6.6) | 8 (3.8) | |
| Neoadjuvant treatment | | | |
| No | 103 (48.8) | 13 (6.2) | |
| Yes | 88 (41.7) | 7 (3.3) | |
| Poorly differentiated carcinoma | | | 0.317 |
| Negative | 129 (61.1) | 16 (7.6) | |
| Positive | 62 (29.4) | 4 (1.9) | |
| T-stage | | | 0.500 |
| pT1 | 52 (24.6) | 4 (1.9) | |
| pT2 | 95 (45.0) | 13 (6.2) | |
| pT3 | 44 (20.9) | 3 (1.4) | |
| pT4 | 0 (.0) | 0 (.0) | |
| N-stage | | | 1 |
| pN0 | 82 (38.9) | 8 (3.8) | |
| pN1 | 109 (51.7) | 12 (5.7) | |
| TNM-stage | | | 0.596 |
| pStage I | 58 (27.5) | 6 (2.8) | |
| pStage II | 63 (29.9) | 4 (1.9) | |
| pStage III | 27 (12.8) | 4 (1.9) | |
| pStage IV | 15 (7.1) | 2 (.9) | |
| Pre-operative CEA | | | 0.576 |
| Within normal | 146 (69.2) | 17 (8.1) | |
| Above normal (>5 U/mL) | 45 (21.3) | 3 (1.4) | |
| Pre-operative CA19-9 | | | 1 |
| Within normal | 99 (46.9) | 10 (4.7) | |
| Above normal (>37 U/mL) | 92 (43.6) | 10 (4.7) | |

Supplementary Table 1 Prevalence of CLDN18.2 positivity by demographic characteristics among all screened patients

Presented as n (%). ASA-PS, American Society of Anesthesiologists Physical Status; CEA, Carcinoembryonic Antigen; CA 19-9, Carbohydrate Antigen 19-9.

| Variables | unmatched | matched | p-value |
|---|---|---|---|
| Overall | 10 (7.5) | 123 (92.5) | |
| Age | | | 0.520 |
| ≤70 | 4 (3.0) | 65 (48.9) | |
| >70 | 6 (4.5) | 58 (43.6) | |
| Sex | | | 0.744 |
| Female | 6 (4.5) | 61 (45.9) | |
| Male | 4 (3.0) | 62 (46.6) | |
| ASA-PS classification system | | | 0.610 |
| 0-2 | 10 (7.5) | 97 (72.9) | |
| 3 | 0 (.0) | 15 (11.3) | |
| BMI | | | 0.106 |
| ≤25 | 6 (4.5) | 92 (69.2) | |
| >25 | 4 (3.0) | 20 (15.0) | |
| Tumor location | | | 0.725 |
| Head | 6 (4.5) | 85 (63.9) | |
| Body/Tail | 4 (3.0) | 38 (28.6) | |
| Resectability | | | 1 |
| R | 7 (5.3) | 82 (61.7) | |
| BR/UR | 3 (2.3) | 41 (30.8) | |
| Neoadjuvant treatment | | | 1 |
| No | 5 (3.8) | 56 (42.1) | |
| Yes | 5 (3.8) | 67 (50.4) | |
| Evans grade | | | 0.559 |
| 1 | 3 (2.3) | 18 (13.5) | |
| 2a/2b | 2 (1.5) | 37 (27.8) | |
| 3 | 0 (.0) | 5 (3.8) | |
| Poorly differentiated carcinoma | | | 1 |
| Negative | 7 (5.3) | 84 (63.2) | |
| Positive | 3 (2.3) | 39 (29.3) | |
| T-stage | | | 0.753 |
| pT1 | 3 (2.3) | 31 (23.3) | |
| pT2 | 4 (3.0) | 63 (47.4) | |
| pT3 | 3 (2.3) | 29 (21.8) | |
| pT4 | 0 (.0) | 0 (.0) | |
| N-stage | | | 0.519 |
| pN0 | 5 (3.8) | 48 (36.1) | |
| pN1 | 5 (3.8) | 75 (56.4) | |
| TNM-stage | | | 0.959 |
| pStage I | 3 (2.3) | 34 (25.6) | |
| pStage II | 4 (3.0) | 44 (33.1) | |
| pStage III | 2 (1.5) | 20 (15.0) | |
| pStage IV | 0 (.0) | 11 (8.3) | |
| Pre-operative CEA | | | 0.690 |
| Within normal | 9 (6.8) | 95 (71.4) | |
| Above normal (>5 U/mL) | 1 (.8) | 28 (21.1) | |
| Pre-operative CA19-9 | | | 0.753 |
| Within normal | 6 (4.5) | 66 (49.6) | |
| Above normal (>37 U/mL) | 4 (3.0) | 57 (42.9) | |

Supplementary Table 2 Prevalence of unmatched CLDN18.2 positivity between biopsy and resected specimens

Presented as n (%). ASA-PS, American Society of Anesthesiologists Physical Status; CEA, Carcinoembryonic Antigen; CA 19-9, Carbohydrate Antigen 19-9.

| Variables | unmatched | matched | p-value |
|---|---|---|---|
| Overall | 10 (16.7) | 50 (83.3) | |
| Age | | | 0.263 |
| ≤70 | 5 (8.3) | 36 (60.0) | |
| >70 | 5 (8.3) | 14 (23.3) | |
| Sex | | | 0.737 |
| Female | 4 (6.7) | 24 (40.0) | |
| Male | 6 (10.0) | 26 (43.3) | |
| ASA-PS classification system | | | 0.190 |
| 0-2 | 7 (11.7) | 42 (70.0) | |
| 3 | 2 (3.3) | 3 (5.0) | |
| BMI | | | 1 |
| ≤25 | 8 (13.3) | 38 (63.3) | |
| >25 | 1 (1.7) | 7 (11.7) | |
| Tumor location | | | 1 |
| Head | 6 (10.0) | 29 (48.3) | |
| Body/Tail | 4 (6.7) | 21 (35.0) | |
| Resectability | | | 0.073 |
| R | 3 (5.0) | 33 (55.0) | |
| BR/UR | 7 (11.7) | 17 (28.3) | |
| Poorly differentiated carcinoma | | | 0.148 |
| Negative | 9 (15.0) | 32 (53.3) | |
| Positive | 1 (1.7) | 18 (30.0) | |
| T-stage | | | 0.356 |
| pT1 | 1 (1.7) | 15 (25.0) | |
| pT2 | 7 (11.7) | 23 (38.3) | |
| pT3 | 2 (3.3) | 12 (20.0) | |
| pT4 | 0 (.0) | 0 (.0) | |
| N-stage | | | 0.305 |
| pN0 | 6 (10.0) | 20 (33.3) | |
| pN1 | 4 (6.7) | 30 (50.0) | |
| TNM-stage | | | 0.664 |
| pStage I | 5 (8.3) | 17 (28.3) | |
| pStage II | 1 (1.7) | 12 (20.0) | |
| pStage III | 1 (1.7) | 9 (15.0) | |
| pStage IV | 2 (3.3) | 6 (10.0) | |
| Post-operative CEA | | | 0.409 |
| Within normal | 7 (11.7) | 40 (66.7) | |
| Above normal (>5 U/mL) | 3 (5.0) | 9 (15.0) | |
| Post-operative CA19-9 | | | 0.577 |
| Within normal | 10 (16.7) | 43 (71.7) | |
| Above normal (>37 U/mL) | 0 (.0) | 6 (10.0) | |
| Adjuvant treatment | | | 0.675 |
| No | 3 (5.0) | 10 (16.7) | |
| Yes | 7 (11.7) | 40 (66.7) | |
| Early recurrence (<1y) | | | 1 |
| No | 5 (8.3) | 30 (50.0) | |
| Yes | 3 (5.0) | 15 (25.0) | |

Supplementary Table 3 Prevalence of unmatched CLDN18.2 positivity between primary and recurrent specimens

Presented as n (%). ASA-PS, American Society of Anesthesiologists Physical Status; CEA, Carcinoembryonic Antigen; CA 19-9, Carbohydrate Antigen 19-9.
